# Decoupling the roles of cell shape and mechanical stress in orienting and cueing epithelia mitosis

**DOI:** 10.1101/177592

**Authors:** Alexander Nestor-Bergmann, Georgina A. Stooke-Vaughan, Georgina K. Goddard, Tobias Starborg, Oliver E. Jensen, Sarah Woolner

## Abstract

Distinct mechanisms involving cell shape and mechanical force are known to influence the rate and orientation of division in cultured cells. However, uncoupling the impact of shape and force in tissues remains challenging. Combining stretching of *Xenopus laevis* tissue with a novel method of inferring relative mechanical stress, we find separate roles for cell shape in orientating division and mechanical stress in cueing division. We demonstrate that division orientation is best predicted by an axis of cell shape defined by the position of tricellular junctions (TCJs), which aligns exactly with the principal axis of local cell stress rather than the tissue-level stress. The alignment of division to cell shape requires functional cadherin and the localisation of the spindle orientation protein, LGN, to TCJs, but is not sensitive to relative cell stress magnitude. In contrast, cell proliferation rate is more directly regulated by mechanical stress, being correlated with relative isotropic stress, and can be decoupled from cell shape when myosin II is depleted.

## INTRODUCTION

Cell division orientation and timing must be carefully regulated in order to shape tissues and determine cell fate, preventing defective embryonic development and diseases such as cancer (Mishra and Chan, 2014; Pease and Tirnauer, 2011; Quyn et al., 2010). Recent work has shown that mechanical cues from the extracellular environment can influence cell division rate (Benham-Pyle et al., 2015; Streichan et al., 2014a) and orientation (Campinho et al., 2013; Legoff et al., 2013; Mao et al., 2011; Fink et al., 2011). What remains unclear is whether dividing cells are directly sensing mechanical forces or are responding to changes in cell shape induced by these forces. This distinction is crucial as the molecular mechanisms involved in either shape-or force-sensing could be very different (Luo et al., 2013; Nestor-Bergmann et al., 2014).

Several mechanisms of division orientation control have been postulated in single cells, with evidence for both shape-and stress-sensing (Fink et al., 2011; Minc et al., 2011; Minc and Piel, 2012; Théry et al., 2006). There is limited understanding of how these models could apply to tissues, where cells are linked together by adhesions and it is far more difficult to exclusively manipulate either cell shape or mechanical stress. Recent evidence for a shape-sensing mechanism was found in the *Drosophila* pupal notum. The spindle orientation protein, Mud (*Drosophila* orthologue of NuMA), localises at tricellular junctions (TCJs), recruiting force generators to orient astral microtubules in rounding mitotic cells (Bosveld et al., 2016). However, this mechanism has yet to be demonstrated in another system or related to mechanical stress. In contrast, recent work in a stretched monolayer of MDCK cells has indicated that division orientation may be mediated by a tension-sensing mechanism requiring E-cadherin, although an additional role for cell shape sensing could not be excluded (Hart et al., 2017). Indeed, divisions in MDCK cells have also been found to align better with cell shape than a global stretch axis, though local cell stress was not known in this case (Wyatt et al., 2015).

Separating the roles of shape and stress in tissues will inevitably require an understanding of how force is distributed through heterogeneous cell layers. Experimental methods of assessing stress include laser ablation, atomic force microscopy and micro-aspiration (Campinho et al., 2013; Davidson et al., 2009; Hoh and Schoenenberger, 1994; Hutson et al., 2003). Whilst informative, these techniques are invasive, perturbing the stress field through the measurement, and usually require constitutive modelling for the measurement to be interpreted (Stooke-Vaughan et al., 2017; Sugimura et al., 2016). However, mathematical modelling combined with high quality fluorescence imaging now provides the possibility of non-invasively inferring mechanical stress in tissues (Brodland et al., 2014; Chiou et al., 2012; Feroze et al., 2015; Ishihara and Sugimura, 2012; Nestor-Bergmann et al., 2017; Xu et al., 2015).

In this article, we apply a reproducible strain to embryonic *Xenopus laevis* tissue to investigate the roles of shape and stress in cell division in a multi-layered tissue. We particularly focus on mathematically characterising local (cell-level) and global (tissue-level) stress and the relation to cell shape and division. Our data suggest that mechanical stress is not directly sensed for orienting the mitotic spindle, acting only to deform cell shape, but is more actively read as a cue for mitosis.

## RESULTS

### Application of tensile force to a multi-layered embryonic tissue

To investigate the relationship between force, cell shape and cell division in a complex tissue, we developed a novel system to apply reproducible mechanical strain to a multi-layered embryonic tissue. Animal cap tissue was dissected from Stage 10 *Xenopus laevis* embryos and cultured on a fibronectin-coated elastomeric PDMS substrate (Figure 1A). A uniaxial stretch was applied to the PDMS substrate using an automated stretch device (Figure 1A), and imaged using standard microscopy. The three-dimensional structure of the stretched tissue (assessed using 3View EM) could be seen to comprise of approximately three cell layers (Figure 1B), as would be expected in a stage 10 *Xenopus laevis* embryo (Keller, 1980; Keller and Schoenwolf, 1977), therefore maintaining the multi-layered tissue structure present *in vivo.*

**Figure 1:**
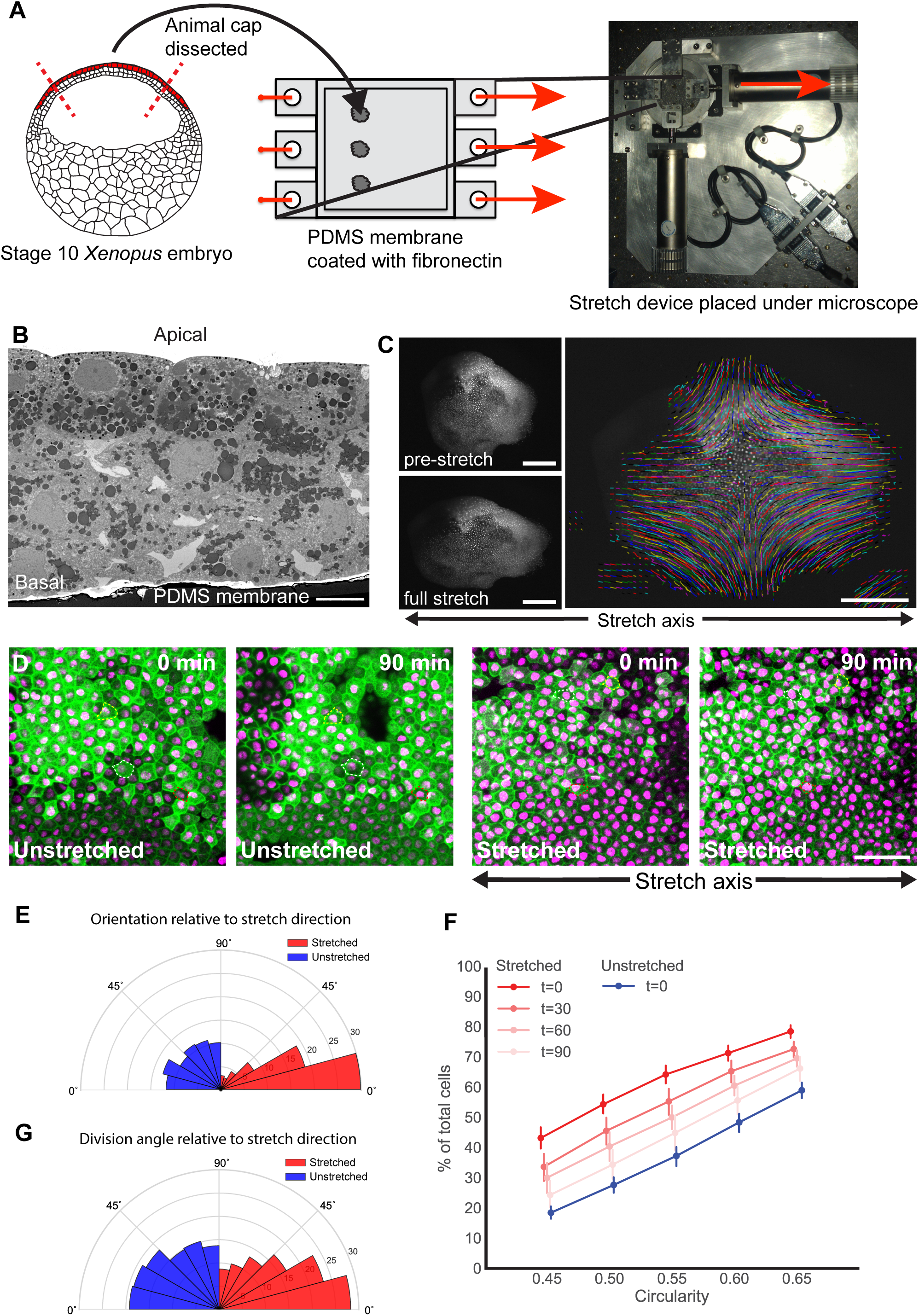
Application of tensile force to a multi-layered tissue. **A.** Animal cap tissue was dissected from Stage 10 *Xenopus laevis* embryos and adhered to fibronectin-coated PDMS membranes and a 35% uniaxial stretch of the membrane was applied. **B**. The animal cap tissue is 2-3 cells thick; cell shape and divisions were assessed in the apical cell layer. **C**. Displacement of nuclei was tracked in a stretched animal cap. **D**. Confocal images of the apical cells in unstretched and stretched animal caps (green: GFP-alpha-tubulin; magenta: cherry-histone2B), taken 0 and 90 minutes after stretch. Representative cells outlined by dashed lines. **E**. Rose plot showing orientation of cell shape relative to direction of stretch in unstretched (blue) and stretched (red; measured immediately following stretch) experiments. **F**. Cumulative plots of cell circularity in unstretched (blue) and stretched (red; at 0, 30, 60 and 90 mins after stretch) animal caps (0=straight line; 1=circle). 100% of cells have circularity ≤ Markers slightly off-set for clarity. **G.** Rose plot of division angle relative to direction of stretch for unstretched (red) and stretched (blue) experiments. Kolmogorov-Smirnov test indicates that the unstretched distribution is not significantly different from a uniform distribution, n = 343 divisions, 15 animal caps; Kolmogorov-Smirnov test indicates that stretched distribution is significantly different from uniform, p < 1.4×10^−9^, n = 552 divisions, 17 animal caps. Scale bar: 10µm in **B**, 500µm in **C**, 50µm in **D**.

### Stretching elongates cell shape and reorients divisions

A 35% stretch of the PDMS substrate led to a 19.67 ± 1.91% (95% confidence interval) elongation of apical cells in the animal cap along the stretch axis (measured change in length of 1-dimensional lines drawn on opposite sides of the animal cap; displacement field shown in Figure 1C). The difference in elongation between the substrate and apical cells is presumably a result of the mechanical stress being dissipated through multiple cell layers. The qualitative change in cell shape was not as substantial as was previously observed in stretched monolayers (Wyatt et al., 2015) (Figure 1D).

We mathematically characterised shape using two parameters: orientation of the principal axis of cell shape relative to the stretch axis (0°), *θ*_*A*_, and cell circularity, C_A_ (derived in Section 1 of the Supplementary Document). C_A_ describes the degree of elongation of a cell (ranging from 0 being a straight line to 1 being a perfect circle) and *θ*_*A*_ indicates the principal direction in which this occurs. Stretching oriented the majority of cells with the direction of stretch (Figure 1E) and caused a highly reproducible elongation of cell shape (Figure 1F). However, when the substrate was held fixed following stretch, cell elongation reduced over time and returned close to the unstretched shape profile after 90 minutes (95% confidence intervals of stretched animal caps at t = 90 minutes overlap with unstretched caps; Figure 1F). Therefore, cells in this tissue adapt to the elongation caused by stretching and do not behave like a purely elastic material.

In unstretched tissue, division orientation, *θ*_*D*_, was not significantly different from a uniform distribution (p = 0.36, Kolmogorov-Smirnov test; Figure 1G). In contrast, divisions in the stretched tissue were significantly oriented along the axis of stretch, (p < 1.43×10^−9^, Kolmogorov-Smirnov test; Figure 1G), with 52% of divisions oriented within 30° of the stretch axis (compared to 36% in unstretched).

### Shape-based models of division differ significantly depending on the cellular characteristics used to define shape

A shape-based ‘long axis’ division rule may explain why stretching reorients divisions. However, the precise molecular mechanism behind shape-based models remains unclear and may vary across cell type and tissue context (Campinho et al., 2013; Fink et al., 2011; Minc et al., 2011). Past models have used different characteristics to determine the shape of a cell, usually selecting one of the following: cell area, cell perimeter and tricellular junction location (TCJ, which we define here as the meeting point of three or more cells). Though often used interchangeably, these shape characteristics model different biological functions. We investigated their differences and determined if one characteristic predicts division orientation better than the others.

We modelled cell shape by area, perimeter and TCJs to derive three respective measures of cell shape orientation, *θ*_*A*_, *θ*_*P*_, and *θ*_*J*_, and circularity, C_A_, C_P_, and C_J_ (Supplementary Document Section 1). Cells tend to have C_P_ > C_A_ > C_J_ i.e. shape generally appears less anisotropic using the perimeter-based measure. C_A_ and C_P_ (and correspondingly *θ*_*A*_ and *θ*_*P*_) are reasonably well correlated, while C_J_ (and *θ*_*J*_) tends to coincide less well with the others (Figure 2A&B and S1A). Thus a cell that appears round by area and perimeter can have clear elongation as measured by TCJs. This is intuitive for rounding mitotic cells, where TCJs can be distributed non-uniformly around the circular periphery (Bosveld et al., 2016). However, it is surprising that this can also be the case in cells with relatively straight edges (Figure 2A’’, note how *θ*_*J*_, – yellow line – differs from *θ*_*A*_ and *θ*_*P*_ – blue and red lines – in the central dark green cells). Notably, cells in the *Xenopus* animal cap do not undergo the dramatic mitotic cell rounding seen in some other systems (Bosveld et al., 2016) (Figure S1B&C).

**Figure 2:**
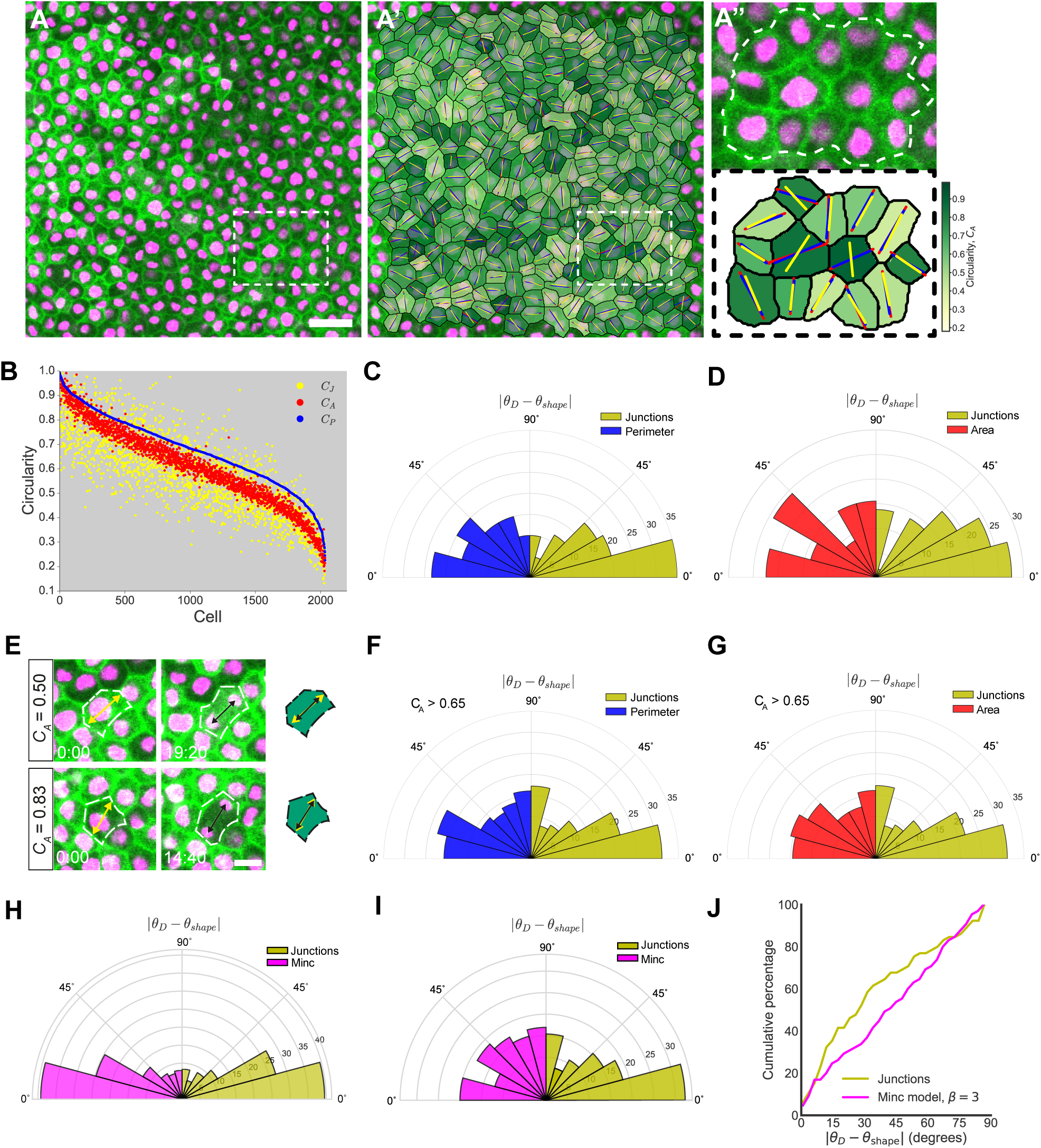
Cell division orientation is best predicted by an axis of shape defined by TCJs. **A.** Representative image of control cells from an unstretched experiment. Scale bar: 20µm **A’.** Overlay of segmentation of cells given in **A**, with the principal axis of shape characterised by area, perimeter and junctions drawn in red, blue and yellow respectively. **A’’.** Enlargement of segmented cells from white box drawn in **A’**, cells analysed are outlined by dashed white line **B.** Circularities of 2035 cells from unstretched experiments, with shape characterised by area, perimeter and junctions plotted in red, blue and yellow respectively. Cells have been ordered in descending order of perimeter-based circularity (C_P_), with the corresponding values of C_A_ and C_J_ plotted alongside. **C**. Rose plot of difference between division angle, *θ*_*D*_, and orientation of shape based on perimeter (blue; *θ*_*shape*_ = *θ*_*P*_) and junctions (yellow; *θ*_*shape*_ = *θ*_*J*_), for cells which satisfy |*θ*_*P*_ − *θ*_*J*_| ≥ 15°. **D**. Rose plot of difference between division angle, *θ*_*D*_, and orientation of shape based on area (red; *θ*_*shape*_ =*θ*_*A*_) and junctions (yellow; *θ*_*shape*_ = *θ*_*J*_), for cells which satisfy |*θ*_*A*_ − *θ*_*J*_| ≥ 15°. **E**. Examples of round (top) and elongated (bottom) cells where division angle (black arrows) is well predicted by the principal axis of shape defined by area (yellow arrows). **F.** Rose plot of difference between division angle, *θ*_*D*_, and orientation of shape based on perimeter (blue; *θ*_*shape*_ = *θ*_*P*_) and junctions (yellow; *θ*_*shape*_ = *θ*_*J*_), for round cells which satisfy C_A_ > 0.65. **G.** Rose plot of difference between division angle, *θ*_*D*_, and orientation of shape based on area (red; *θ*_*shape*_ =*θ*_*A*_) and junctions (yellow; *θ*_*shape*_ = *θ*_*J*_), for round cells which satisfy C_A_> 0.65. See also Figure S1. **H** Rose plot of difference between division angle, *θ*_*D*_, and orientation of shape based on Minc model when β = 3 (magenta; *θ*_*shape*_ = *θ*_*Minc*_) and junctions (yellow; *θ*_*shape*_ = *θ*_*J*_) for all cells in stretched and unstretched experiments (n = 599 cells). **I.** Rose plot of difference between division angle, *θ*_*D*_, and orientation of shape based on Minc model when β = 3 (magenta; *θ*_*shape*_ = *θ*_*Minc*_) and junctions (yellow; *θ*_*shape*_ = *θ*_*J*_), for cells which satisfy |*θ*_*Minc*_ − *θ*_*J*_| ≥ 15° (n = 65 cells). **J.** Cumulative plot of difference between division angle, *θ*_*D*_, and orientation of shape for data shown in I.

### TCJ placement is a better predictor of division orientation than cell area, cell perimeter or microtubule length

Given that *θ*_*A*_, *θ*_*P*_, and *θ*_*J*_ are often highly correlated, division orientation is generally well predicted by all three. We therefore focused on cases in which the orientations of shape differed by at least 15°. In a pooled sample of 600 cells from stretched and unstretched tissue, only 7 cells were found to have | *θ*_*A*_ − *θ*_*P*_| ≥ 15°. 58 satisfied |*θ*_*A*_ − *θ*_*J*_| ≥ 15° and 60 satisfied |*θ*_*P*_ − *θ*_*J*_| ≥ 15°. In the latter two cases, **θ**_*J*_ was a significantly better predictor of division angle than random (p < 0.0162 when |*θ*_*A*_ − *θ*_*J*_| ≥ 15° p < 0.0042 when |*θ*_*P*_ − *θ*_*J*_| ≥ 15° Mann-Whitney U test), but *θ*_*A*_ and *θ*_*P*_ were not (Figure 2C&D). Furthermore, C_A_, C_P_, and C_J_ were all significantly higher in these subpopulations (Figure S1D&E; 95% confidence intervals do not overlap), indicating that these cells are rounder, yet can still effectively orient their spindle in-line with their TCJs. This result is strengthened considering that TCJs provide fewer data points than area or perimeter, thus junctional data may be more susceptible to geometric error than area and perimeter. For all of our data comparing cell shape with division orientation, we use shape determined just prior to nuclear envelope breakdown (NEB), avoiding any possible shape changes due to mitosis (e.g. cell rounding on entry into mitosis/elongation at anaphase). However, to test whether the fidelity of division alignment to TCJ shape changes depending on when shape is measured we compared |*θ*_*D*_ *- *θ**_*J*_| at time-points through mitosis, finding no significant difference (Figure S1F). It is important to note that we do not see significant cell rounding in the *Xenopus* animal cap upon entry into mitosis (Figure S1C), so static fidelity is likely a reflection of relatively static cell shape in this system, a feature which helps simplify our analysis.

In unstretched tissue, cells which we classed as “rounded” (C_A_ > 0.65; Figure 2E) showed no significant correlation between *θ*_*A*_ and *θ*_*D*_ or *θ*_*P*_ and *θ*_*D*_, as could be expected from previous work (Minc et al., 2011). However, *θ*_*J*_ was significantly aligned with division angle in these round cells, when compared to random (p = 0.025, Mann-Whitney U test) (Figure 2F&G). This degree of sensitivity is striking and further demonstrates that TCJ-sensing could function effectively in round cells, which may have previously been thought to divide at random.

Our analysis is based purely on predictions arising from the data and thereby has the advantage of being independent of unknown model parameters and assumptions. However, to test how our division predictions compare with previous models of division orientation we turned to a well-known shape-based model of division in isolated cells (Minc et al., 2011). The “Minc” model hypothesises that astral microtubules exert length-dependent pulling forces on the spindle, thereby exerting a torque and rotating the spindle, with division predicted to occur along the axis of minimum torque. In this shape-based model, the shape of the cell determines the distribution of torque on the spindle and thereby the division axis (see the Supplementary Document for further details of this model and its implementation). As with our purely geometric measures of shape, we found that the Minc model predicts division orientation significantly better than a random distribution (Figure 2H; p < 4.1×10^−40^ for TCJs and p < 1.2×10^−39^, Mann-Whitney U test). However, for cells where the predicted division axes according to TCJs (*θ*_*J*_) and the Minc model (*θ*_*Minc*_) differed by more than 15°, TCJs (*θ*_*J*_) provided a prediction of division angle that was significantly better than random (p < 0.028, Mann-Whitney U test), while division predicted by microtubule pulling forces (*θ*_*Minc*_) did not (Figure 2I & J), indicating that TCJs provide a better prediction of division orientation. This result held for multiple scaling laws between microtubule length and force (Figure S1G&H).

### Local cell shape aligns with local stress and predicts division orientation better than global stretch and stress

Contrary to observations in monolayers (Hart et al., 2017), we found that cells in stretched tissue divide according to cell shape both when *θ*_*J*_ is oriented with (Figure 3A) and against (Figure 3B&C) the direction of stretch. Moreover, in the case of cells that are relatively round in shape (C_J_>0.65), there is no preference for aligning with the global stretch direction and indeed alignment with TCJ shape still appears more accurate than with the stretch axis (Figure S2A&B; p < 0.005 for TCJS, not significant for stretch direction, Mann-Whitney U test). These data indicate that global stretch direction is a poor predictor of division angle when compared to cell shape. However, little is known about the local stress distribution around individual cells in a tissue subjected to a stretch, which may not coincide with global stress in such a geometrically heterogeneous material.

**Figure 3:**
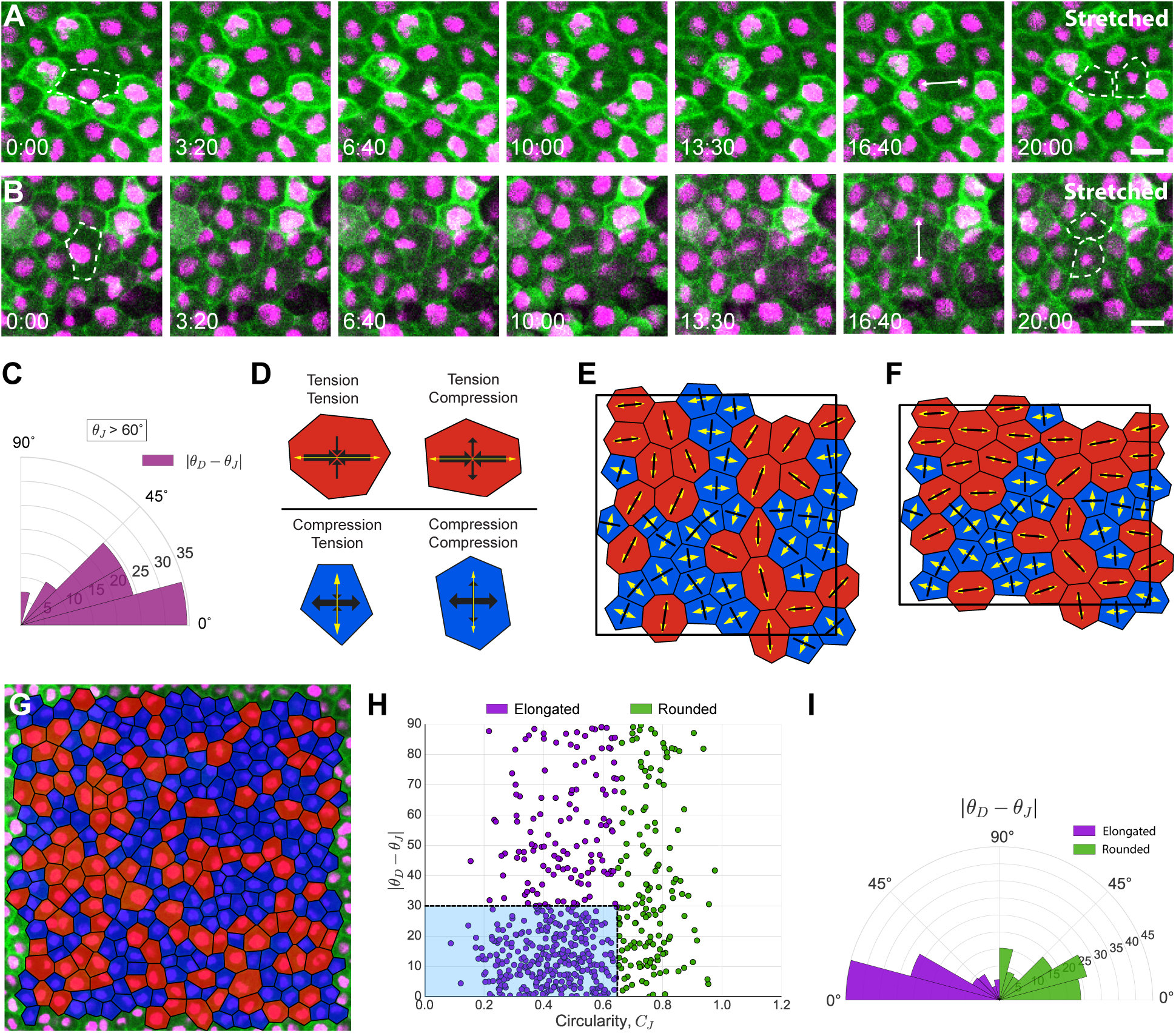
Division orientation is better predicted by shape in elongated cells, rather than those with higher relative isotropic or shear stress. **A.** Images taken from a confocal timelapse movie of a division in a cell in stretched tissue whose interphase shape (dashed line, 0:00) is oriented with the stretch (horizontal) axis. Cell division aligns with both cell shape and stretch axis. **B.** Timelapse images of an unusual cell in a stretched tissue, whose interphase shape (dashed line, 0:00) is oriented against the stretch axis. Cell division aligns with cell shape but against the stretch axis. **C.** Rose plot of difference between division angle, *θ*_*D*_, and orientation of shape based on junctions, *θ*_*J*_, for cells from stretched experiments, where *θ*_*J*_ was at least 60 °divergent to the direction of stretch. 29 cells satisfied this condition. Kolmogorov-Smirnov test found a significant difference from a uniform distribution (p=0.022). **D.** Representative cells showing classification of cell stress configurations. Red (blue) cells are under net tension (compression), where *p*^eff^ is positive (negative). Larger (smaller) black arrows indicate the orientation of the principal (secondary) axis of stress, with inward-(outward)-pointing arrows indicating the tension (compression) generated by the cell. Yellow arrows indicate the principal axis of shape defined by cell junctions, which aligns exactly with a principal axis of stress. **E.** 50 simulated cells randomly generated in a periodic box, relaxed to equilibrium with parameters (Λ,Г = (−0.259, 0.172), under conditions of zero global stress (Nestor-Bergmann et al., 2017). Red (blue) cells are under net tension (compression). Principal axis of stress (shape) indicated in black (yellow). **F.** Cells from **E** following a 13% area-preserving uniaxial stretch along the x-axis. **G.** Example segmented cells from an unstretched experiment. Cells in red (blue) are predicted to be under net tension (compression). **H.** Cell circularity defined by junctions, C_J_, vs |*θ*_*D*_ − *θ* _*J*_| Spearman rank correlation coefficient found a significant correlation (p < 3.04 × 10^−10^). Elongated cells (C_J_≤0.65) cluster in blue box, whereas rounded cells (C_J_ >0.65) have a more uniform distribution. **I.** Rose plot of difference between division angle, *θ*_*D*_, and orientation of shape based on junctions, *θ*_*J*_ for round (C_J_>0.65; right) and elongated (C_J_≤0.65; left) cells shown in **H**. Mann-Whitney U test indicated that elongated cells have *θ*_*J*_ aligned significantly more with *θ*_*D*_ than rounded cells (p < 1.64 × 10^−8^). Scale bar in **A**&**B:** 20μm. All rose plots show percentage of cells.

We extended a popular vertex-based model to mathematically characterise cell stress (Brodland et al., 2014; Chiou et al., 2012; Ishihara and Sugimura, 2012; Nestor-Bergmann et al., 2017; Nestor-Bergmann et al., 2018). Predicted orientations of forces from the model have been found to be in accordance with laser ablation experiments (Farhadifar et al., 2007; Landsberg et al., 2009), indicating that the model can provide a physically relevant description of cellular stresses. Our methodology allows relative cell stress to be inferred solely from the positions of cell vertices, without invasively altering the mechanical environment (Supplementary Document, Section 2). The model predicts that the orientation of cell shape based on TCJs, *θ*_*J*_, aligns exactly with the principal axis of local stress (Nestor-Bergmann et al., 2017) (Figure 3D). We demonstrated this computationally in stretched tissue by simulating a uniaxial stretch (Figure 3E-F). Following stretch, we see that local cell stress remains aligned with *θ*_*J*_, rather than the global stress along the x-axis. Much previous work assumes that the local axis of stress coincides with the global stress. Significantly, the model predicts that a stress-sensing mechanism would align divisions in the same direction as a shape-based mechanism (as in Figure 3B).

### The magnitude of cell stress does not correlate with the alignment of division angle and TCJ positioning

If a stress-sensing mechanism were contributing to orienting division, we hypothesised that cells under higher net tension or compression might orient division more accurately with the principal axis of stress (*θ*_*J*_). We infer relative tension/compression using the isotropic component of stress, effective pressure (*P*^eff^) (Nestor-Bergmann et al., 2017):

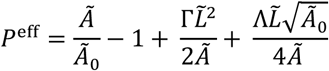

where Ã is cell area, 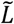 is perimeter, *Ã*_0_ is the preferred area and (Λ, Γ) are model parameters, defined in Section 2 of the Supplementary Document and inferred from data (Nestor-Bergmann et al., 2017). Cells under net tension have *P*^eff^ > 0, whereas *P*^eff^ < 0 indicates net compression. We provide a novel method for estimating *Ã*_0_ in Section 3 of the Supplementary Document. A representative segmentation, showing cells predicted to be under net tension and compression, from an unstretched experiment is given in Figure 3G. Interestingly, we found no correlation between the value of *P*^eff^ (relative isotropic stress) and the alignment of division orientation to *θ*_*J*_ (|*θ*_*D*_ − *θ*_*J*_|) (Figure S2C). The mechanical state of a cell may also be characterised by shear stress, ξ (defined as the eigenvalue of the deviatoric component of the stress tensor, see Section 2 of the Supplementary Document). Larger values of |ξ| indicate increased cellular shear stress. Again, we found no correlation between ξ and the alignment of division to *θ*_*J*_ (Figure S2D).

Despite the lack of correlation with stress magnitude, cell shape anisotropy, measured by C_J_, correlates significantly with |*θ*_*D*_ − *θ*_*J*_| (p < 3.04×10^−10^, Spearman rank correlation coefficient; Figure 3H), with elongated cells having *θ*_*D*_ aligned with *θ*_*J*_ significantly better than round cells (p < 1.64 × 10^−8^; Figure 3I).

### Cadherin is required for positioning the mitotic spindle relative to cell shape

Immunofluorescence staining of β-catenin confirmed that adherens junctions were distributed along the apical cell cortex, but particularly concentrated at the meeting points of three or more cells (Figure 4E). To test a functional requirement for adherens junctions in orienting the spindle, we focused on maternal C-cadherin (cadherin 3), which is expressed at the highest level in Stage 10-11 *Xenopus* embryos (Heasman et al., 1994; Lee and Gumbiner, 1995). We used two constructs to manipulate C-cadherin in the tissue: C-cadherin FL-6xmyc (CdhFL: Full length C-cadherin with 6xmyc tags at the intracellular c-terminus) and C-cadherin ΔC - 6xmyc (CdhΔC: C-cadherin with extracellular and transmembrane domains, but lacking the cytosolic domain) (Figure 4A) (Kurth et al., 1999). CdhFL- and CdhΔC-injected embryos developed normally up to Stage 10/11 (Figure S3A), but the majority of embryos failed to complete gastrulation (Lee and Gumbiner, 1995) (and data not shown). We observed no change in the cumulative distribution of cell circularities in CdhFL-and CdhΔC-injected tissues compared to control tissue (Figure S3B). We also saw no difference in the rate of cell divisions (data not shown).

**Figure 4:**
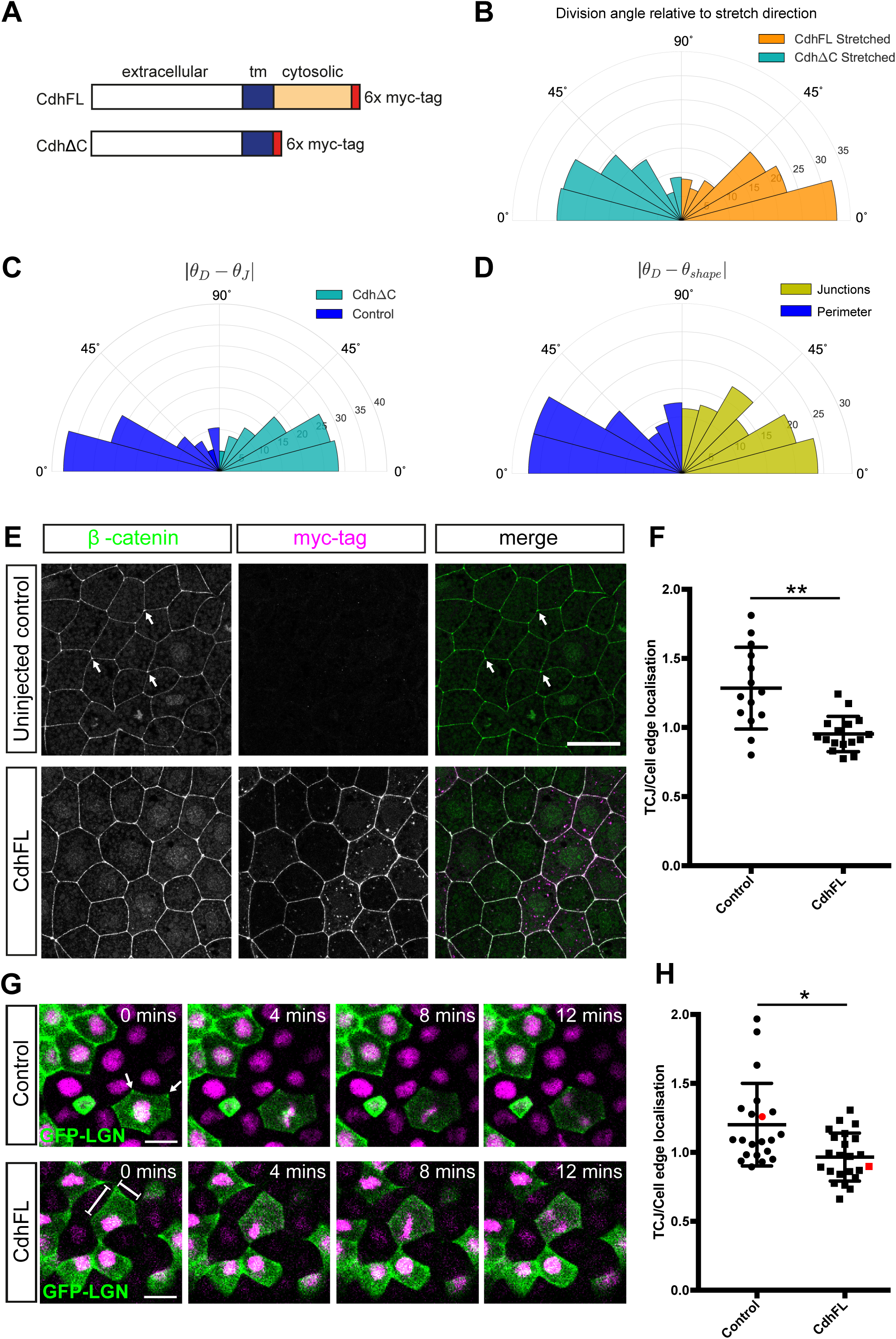
C-Cadherin is required to orient the mitotic spindle according to cell shape. **A.** Schematic of Cadherin contructs CdhFL and CdhΔC **B.** Rose plot of division angles, *θ*_*D*_, relative to direction of stretch for cells from stretched CdhΔC-injected (411 cells; cyan) and stretched CdhFL-injected experiments (552 cells; orange). CdhFL-injected cells align significantly better with direction of stretch (p < 0.0162, Mann-Whitney U test). **C.** Rose plot of difference between division angle, *θ*_*D*_, and orientation of shape based on junctions, *θ*_*J*_, for cells from CdhΔC-injected experiments (390 cells; cyan) and control experiments (239 cells; blue). Distributions are significantly different (p < 0.016 Kolmogorov-Smirnov test). **D.** Rose plot of difference between division angle, *θ*_*D*_, and orientation of shape based on perimeter, *θ*_*P*_, (blue) and junctions, *θ*_*J*_, (yellow) for 96 cells from CdhFL-injected experiments which satisfied |*θ*_*P*_ − *θ*_*J*_| ≥ 15°. *θ*_*D*_ aligns significantly better to *θ*_*P*_ than a random distribution (p < 0.004; Kolmogorov-Smirnov test), but not to *θ*_*J*_. **E**. Single confocal slices from immunofluorescent staining for β-catenin (green) and myc-tag (magenta) in uninjected and CdhFL-injected St 12 embryos (stage matched to time that animal caps are stretched and imaged). Hotspots of β-catenin localization (arrows) are seen at TCJs in controls but are lost when CdhFL is overexpressed. **F.** Quantification of β-catenin-GFP localization at TCJs compared to cell edges in single mitotic cells in animal caps. β-catenin-GFP is more strongly localised at TCJs compared to cell edges in controls but this bias is lost in CdhFL-injected tissue (*p < 0.01 Komogorov-Smirnov test; n = 14, 16 mitotic cells from 6, 4 unrelated animal caps for control and CdhFL, respectively). **G**. Time-lapse movies of control and CdhFL-injected animal cap tissue expressing GFP-LGN in a mosaic fashion. In control cells, GFP-LGN is enriched at TCJs during interphase (arrows) and this localization persists through mitosis. The enrichment of GFP-LGN at TCJs is lost when CdhFL is expressed, with localisation spread throughout the cell edge. **H.** Quantification of GFP-LGN localisation at TCJs compared to cell edges in single mitotic cells in animal caps. GFP-LGN is more strongly localised at TCJs compared to cells edges in controls but this bias is lost in CdhFL-injected tissue (*p < 0.05 Komogorov-Smirnov test; n = 21, 23 mitotic cells from 7, 6 unrelated animal caps for control and CdhFL, respectively). Red points show quantification for mitotic cells highlighted in G. Rose plots show percentage of cells. Scale bars represent 20µm. See also Figure S3.

CdhΔC-injected tissue was elongated by application of stretch (Figure S3C), but showed a worse alignment of divisions to stretch direction compared to uninjected control and CdhFL-injected tissue (Figure 4B; Mann-Whitney U test p < 0.0162 for CdhΔC less than CdhFL). Moreover, unstretched CdhΔC-injected tissue showed a significant decrease in the alignment of division angle to *θ*_*J*_, when compared to uninjected controls (Figure 4C; p < 0.016 Kolmogorov-Smirnov test on distributions differing), though both were significantly different to random (control: p < 3.6×10^−11^; CdhΔC: p < 4.3×10^−11^; Kolmogorov Smirnov test). To further investigate a requirement for adherens junctions in division orientation, we overexpressed C-cadherin in the cell cortex by injecting CdhFL. Focussing on cells which satisfied |*θ*_*P*_ − *θ*_*J*_| ≥ 15°, we found the striking result that division orientation was now significantly well predicted by cell perimeter, but no longer by TCJs (Figure 4D; p < 0.0027 for alignment *θ*_*D*_ to *θ*_*P*_, but not significant for *θ*_*D*_ to *θ*_*J*_; Mann-Whitney U test). Therefore, overexpression of CdhFL was sufficient to switch division orientation from alignment with TCJs to alignment with the shape of the entire cortex.

To investigate the mechanism behind the observed switch in division orientation, we explored how overexpression of CdhFL alters the localisation of spindle orientation machinery at the cell cortex. We found that overexpression of CdhFL led to a loss of the “hotspots” of β-catenin localisation at TCJs seen in control tissue, in both interphase and mitotic cells (Figure 4E and F). When CdhFL is overexpressed, β-catenin is more equally spread around the entire apical perimeter of the cell (Figure 4E and F). The “hotspots” of β-catenin localisation in controls are not purely a result of more cells contributing to this focal point, but are also seen when fluorescence intensity is measured in single β-catenin-GFP expressing cells in the animal cap (Figure 4F). To determine how this observed change in adherens junction localisation might alter spindle orientation we investigated how the localisation of the spindle orientation protein, LGN, was altered by overexpression of CdhFL. Mosaic expression of GFP-LGN allowed us to analyse at the single-cell level in stretched and unstretched animal caps. In control tissue LGN, like β-catenin, shows a more concentrated localisation at TCJs (Figure 4G and H). We observed no significant difference in LGN localisation between unstretched and stretched tissue (data not shown). However, we saw a loss of concentrated “hotspots” of LGN localisation when CdhFL is overexpressed, with LGN instead spread more equally around the whole perimeter (Figure 4G and H). We conclude that overexpression of CdhFL switches division orientation from alignment with TCJs to alignment with the shape of the whole cortex by altering the localisation of LGN. Moreover, our data indicate that, in normal circumstances, the concentration of LGN at TCJs plays an important role in aligning divisions with cell shape.

### Cell division rate is temporarily increased following change in global stress

Stretch elicited a reproducible and significant increase in cell division rate, with 6.47 ± 1.12 % of cells dividing per hour in the stretched tissue compared to 3.22 ± 0.55 % in unstretched tissue (Figure 5A, 95% confidence intervals do not overlap), as reported for cultured cells and monolayers (Fink et al., 2011; Streichan et al., 2014b; Wyatt et al., 2015). We roughly classify two distinct periods of division after stretch; there is an initial period of high proliferation (8.1% cells undergoing division per hour; Figure 5B), which drops, after 40-60 minutes, to near-unstretched control levels (4.2% cells undergoing division per hour). Stretching increases apical tissue area by 6 ± 2.69% (95% confidence interval), and is predicted to increase global stress by increasing individual values of *P*^eff^. We sought to determine whether the increase in division rate is a response to these changes.

**Figure 5:**
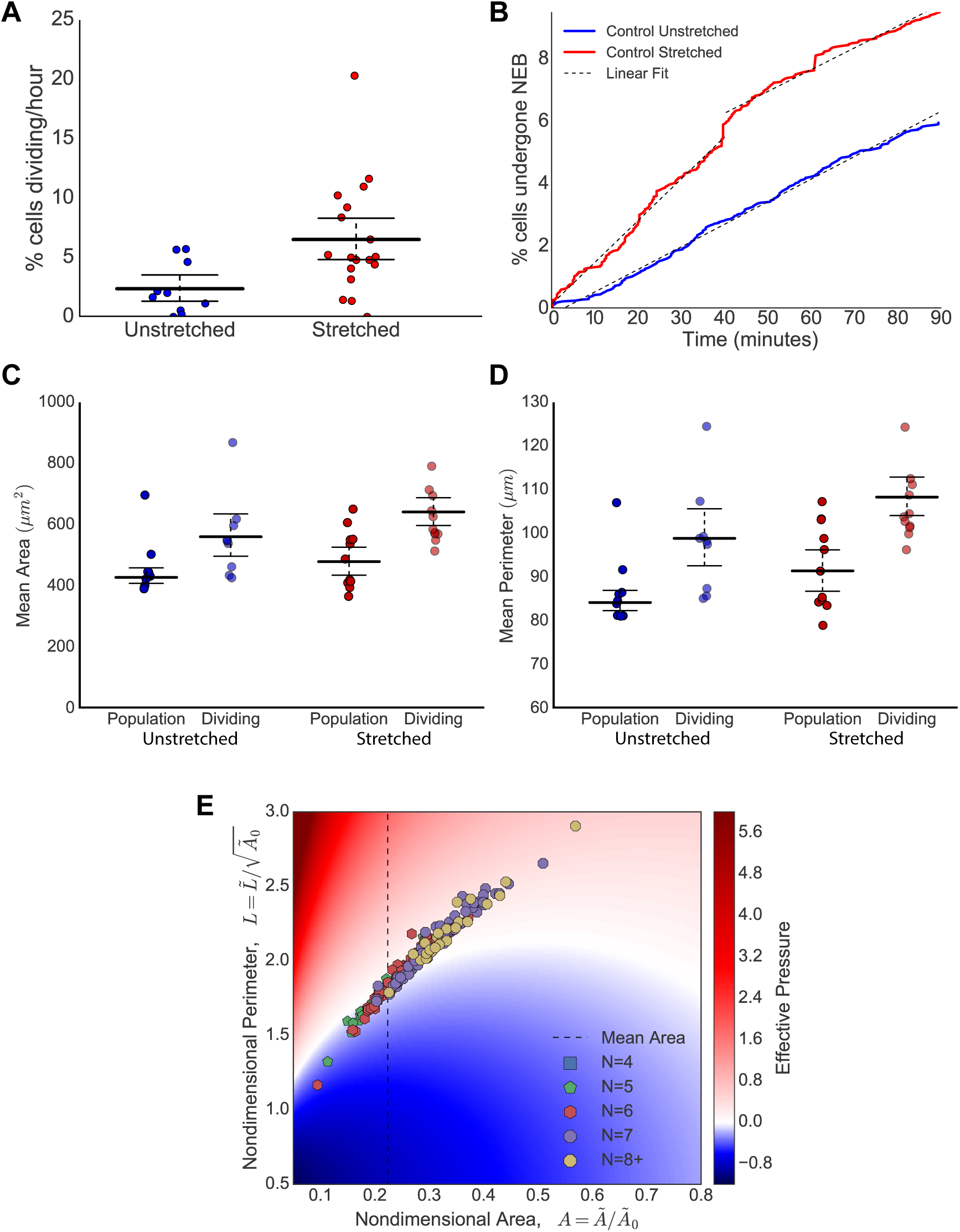
Stretching increases division rate. Dividing cells have large area, perimeter and relative effective pressure. **A.** Division rate (percentage of cells entering mitosis per hour) increases in stretched tissue compared to unstretched. 95% confidence intervals do not overlap, indicating significant difference. Each point represents the mean division rate from an animal cap. **B.** Percentage of cells that have undergone nuclear envelope breakdown (NEB) with respect to time in control stretched (red) and unstretched (blue) experiments from **A**. Dashed lines indicate linear lines of best fit; control unstretched experiments have gradient 4.2% cells undergoing division per hour. Stretched experiments have initial gradient 8.1% and then 4.35% cells undergoing division per hour. **C**. Comparison of mean area of population of all cells vs dividing cells from unstretched and stretched control experiments. Error bars represent mean and 95% confidence intervals, which do not overlap between the population and dividing cells, indicating a significant difference. **D**. Comparison of mean perimeter of population of all cells vs dividing cells from unstretched and stretched control experiments. Error bars represent mean and 95% confidence intervals, which do not overlap between the population and dividing cells, indicating a significant difference. E. Heat map showing predicted relative isotropic stress (effective pressure, *P*^eff^) of dividing cells from control unstretched experiments. Areas and perimeters have been nondimensionalised using the preferred areas, *Ã*_0_, fitted to each experiment in Supplemental Figure 4C. Polygonal class (number of neighbours) indicated by marker colour and style, with (4,5,6,7,8+) sided cells given in (blue, green, red, purple, yellow). Dashed vertical line represents mean area of all cells. Cells lying in red (blue) regions are under predicted net tension (compression).

In both stretched and unstretched experiments, dividing cells had a larger area than the population, being about 22.7% and 25.7% larger on average respectively (Figure 5C). Similarly, the mean perimeter was significantly larger in the dividing cells by about 14.1% in unstretched and 13.8% in stretched (Figure 5D). However, there was no significant difference in the level of cell elongation in dividing cells (Figure S2E). Crucially, we found that dividing cells were more likely to be under predicted net tension than compression (Figure 5E, more cells in red region). However, *P*^eff^ is correlated with cell area (though the two are not always equivalent), thus a further perturbation was required to separate their effects.

### Loss of myosin II reduces cell contractility

We perturbed the mechanical properties of the tissue with targeted knockdown of non-muscle myosin II using a previously published morpholino (Skoglund et al., 2008). As expected, myosin II knockdown disrupted cytokinesis, seen by the formation of ‘butterfly’ shaped nuclei, where daughter cells had not fully separated (Figure 6A&B). However, division rate and orientation could still be assessed using the same methods described for control tissue. Myosin II is known to generate contractility within a tissue (Clark et al., 2014; Effler et al., 2006; Gutzman et al., 2015). Accordingly, we found evidence for reduced contractility in the myosin II MO tissue by observing that cells were much slower at adapting to stretch, remaining elongated for longer (compare Figure 6C to Figure 1F).

**Figure 6:**
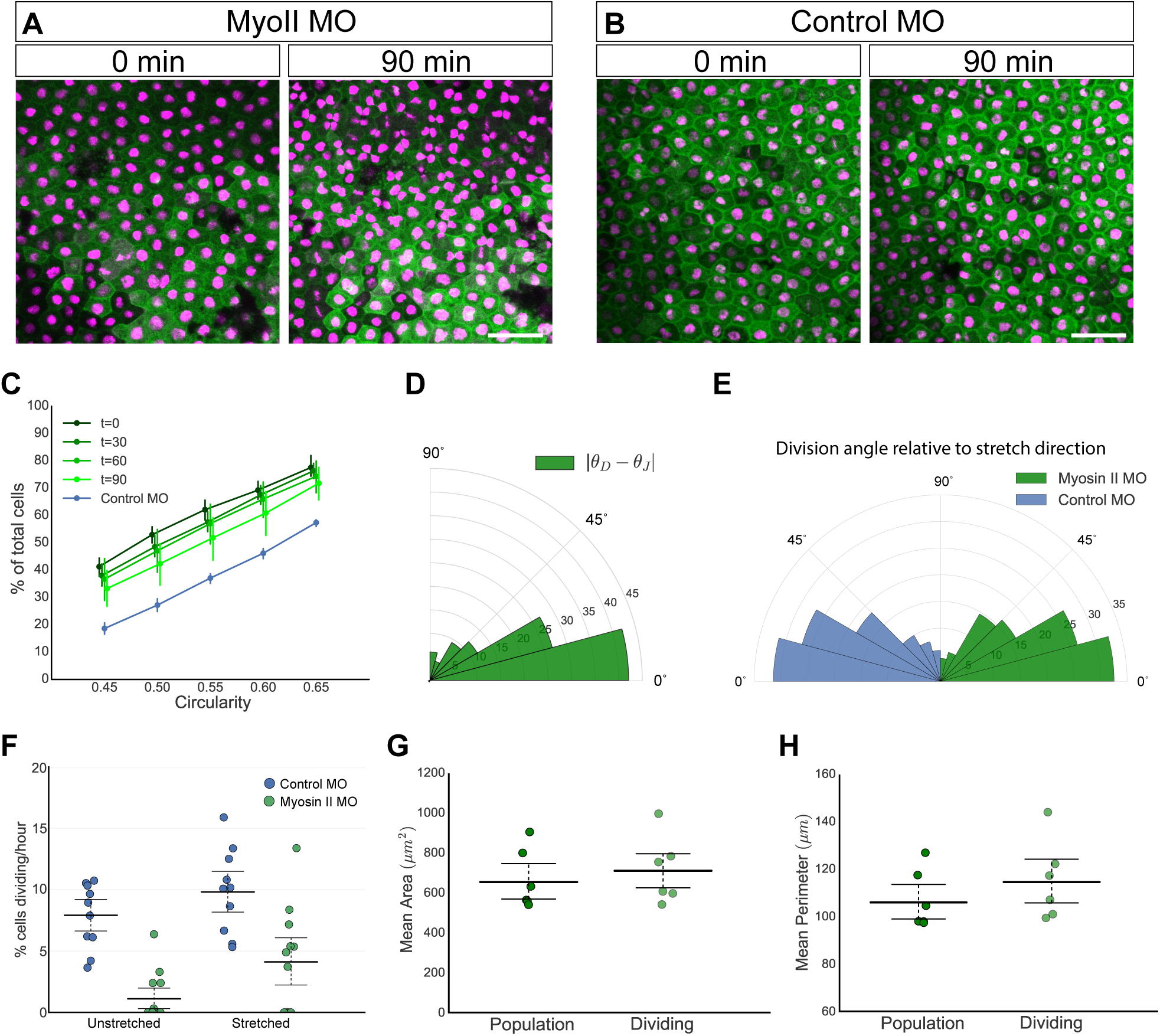
Myosin II MO cells maintain alignment of division to TCJ shape, but have perturbed proliferation rate. **A.** Images taken from a confocal timelapse movie of stretched myosin II morpholino injected animal cap explants at 0 and 90 minute intervals. Butterfly nuclei seen prominently at 90 minutes, where nuclei are in contact. **B.** Timelapse images of control morpholino-injected stretched animal cap explants at 0 and 90 minute intervals. **C.** Cumulative distribution of cell circularity defined by area, C_A_, in myosin II MO knockdown stretched animal caps (shaded green) at t=0, 30, 60 and 90 minutes after stretch. Cumulative distribution for unstretched t=0 control MO knockdown experiments shown in blue. Error bars represent 95% confidence intervals. Error bars for myosin II MO t=90 minutes distribution does not overlap with control MO, indicating a significant difference from unstretched shape. Markers are slightly off-set for clarity. **D.** Rose plot of difference between division angle, *θ*_*D*_, and orientation of shape based on junctions, *θ*_*J*_, for 216 cells from myosin II knockdown stretched experiments. Mann-Whitney U test found significant alignment compared to random (p < 5.72×10^−15^), but no significant difference from equivalent dataset in control stretched experiments. Percentages of cells shown. **E**. Rose plot of division angle relative to direction for stretch for control MO (532 cells; blue) and myosin II MO (301 cells; green) experiments. Mann-Whitney U and Kolmogorov Smirnov test found no significant difference between the two. **F.** Division rate (percentage of total cells entering mitosis per hour) in unstretched and stretched tissue from myosin II MO (green; n=10 for unstretched and n=12 for stretched) and control MO (blue; n=13 for unstretched and n=10 for stretched) experiments. Error bars represent mean and 95% confidence intervals. **G.** Comparison of mean area of population of all cells vs dividing cells from stretched myosin II knockdown experiments. Error bars represent mean and 95% confidence intervals, which overlap, indicating no significant difference. **H.** Comparison of mean perimeter of population of all cells vs dividing cells from stretched myosin II knockdown experiments. Error bars represent mean and 95% confidence intervals, which overlap, indicating no significant difference. Scale bar in **A** and **B**: 100µm.

### Myosin II is required for mitotic entry in unstretched tissue

Somewhat surprisingly, considering suggestions that myosin II may play a stress-sensing role in orienting the spindle (Campinho et al., 2013), we found that alignment of division angle to stretch and *θ*_*J*_ was unaffected in global myosin II knockdown experiments (Figure 6D&E). In contrast, proliferation rate was significantly affected, with divisions virtually ceasing in unstretched myosin II MO tissue. Strikingly, stretching the myosin II MO tissue increased the division rate to significantly higher levels (Figure 6F). Thus myosin II is required to cue cells into division in the unstretched tissue, but this can be partially overridden by applying an external loading. Unlike in control experiments, dividing cells in myosin II knockdown stretch experiments were not significantly larger than the population in area (Figure 6G) or perimeter (Figure 6H), thus cell area has been uncoupled as a cue to divide in the myosin II knockdowns. This finding, along with our observation that dividing cells were more likely to be under relative net tension than relative compression (Figure 5E), indicates that in a tissue the cue to divide, in contrast to division orientation, is directly sensitive to mechanical force.

## DISCUSSION

Previous models of cell division have demonstrated that specific features of cell shape, such as the cell cortex or TCJs, may be important in orienting the spindle (Bosveld et al., 2016; Hertwig, 1893; Luxenburg et al., 2011; Minc et al., 2011). We have presented a framework for characterising cell shape in terms of its area, perimeter or TCJs (Supplementary Document). We find that the principal axis of shape defined by TCJs is the best predictor of division angle, better than cell shape as determined by area, perimeter or by a previous shape-sensing model based on microtubule length (Minc et al., 2011). Moreover, the principal axis of shape defined by TCJs aligns exactly with the principal axis of local stress (Nestor-Bergmann et al., 2017), providing a non-invasive way to infer mechanical stress in individual cells in the epithelium. However, division angle is not better predicted in cells with higher/lower relative isotropic or shear stress, suggesting that cell-level mechanical stress is not a direct cue to orient the spindle. Our findings share similarities with observations in the *Drosophila* pupal notum, where TCJs have been hypothesised to localise force generators to orient the spindle (Bosveld et al., 2016). Notably, however, *Xenopus* animal cap cells do not undergo the dramatic mitotic rounding exhibited by cells in the notum.

Cell-cell adhesion has been linked to spindle orientation in MDCK cells, where E-cadherin instructs LGN/NuMA assembly at cell-cell contacts to orient divisions (Gloerich et al., 2017). E-cadherin polarises along a stretch axis, reorienting divisions along this axis rather than according to cell shape (Hart et al., 2017). In accordance, we find division is less well predicted by shape in embryos injected with C-cadherin ΔC-6xmyc, lacking the cytosolic domain. Interestingly, over-expression of C-cadherin around the entire cell cortex leads to a switch in division orientation, from TCJs to division best predicted by a perimeter-based shape axis. As β-catenin is increased around the cell cortex when C-cadherin is overexpressed, we hypothesized that this may lead to altered recruitment of spindle orientation proteins, such as LGN and NuMA (Gloerich et al., 2017). Indeed we find that while LGN is normally most highly localised to TCJs, overexpression of C-cadherin leads to a loss of these “hotspots” and instead a more even spread of LGN around the entire cell perimeter. We suggest that in the wild type situation, the hotspots of LGN localisation at TCJs will recruit more NuMA and dynein providing localised force generation to orient the spindle according to TCJ shape. When C-cadherin is overexpressed, and the LGN hotspots are no longer present, we suggest that a perimeter-based shape sensing mechanism, similar to that proposed by Minc et al (Minc et al., 2011) predominates. Our TCJ-based system of spindle orientation is similar to the Mud-dependent TCJ-sensing mechanism in the *Drosophila* pupal notum (Bosveld et al., 2016). However, it is important to note a key difference: NuMA, the vertebrate homologue of *Drosophila* Mud, localises to the nucleus during interphase, only localising to the cortex after NEB (Bowman et al., 2006; Kiyomitsu et al., 2012; Seldin et al., 2013; Gloerich et al., 2017). Of future interest will be to determine in vertebrate tissue how the TCJ localisation of LGN influences the highly dynamic recruitment of NuMA. Indeed, recent work in MDCK cells has shown that on mitotic entry, when NuMA is released from the nucleus, it competes LGN away from E-cadherin at the cortex to locally form the LGN/NuMA complex (Gloerich et al., 2017); it will be important to determine if this is happening specifically at TCJs.

Stretching increases proliferation rate, which correlates with cell area, perimeter and effective pressure. We see almost no proliferation in unstretched myosin II MO experiments, although, rather strikingly, the division rate is significantly increased following stretch. Dividing myosin II MO cells are not significantly larger in area or perimeter than the population as a whole, indicating that cell area has been decoupled as a division cue. Considering the established role of myosin II as a force generator (Clark et al., 2014; Gutzman et al., 2015; Vicente-Manzanares et al., 2009), it is possible that the myosin II MO cells cannot generate enough internal contractility in neighbouring cells to engage the mechanical cues required for mitotic entry. Myosin II has also been shown to function in stress-sensing pathways (Hirata et al., 2015; Priya et al., 2015), which may explain why the proliferation rate in stretched myosin II MO cells does not reach the levels of stretched controls. Contrary to findings in other systems (Campinho et al., 2013), a global loss of myosin II does not alter division orientation relative to cell shape. However, future work should look to explore whether anisotropic biases in junctional myosin II affect division orientation, as was recently seen in the Drosophila germband (Scarpa et al., 2018).

In conclusion, we have combined whole-tissue stretching with a biomechanical model to propose separate roles for cell shape and mechanical stress in orienting the spindle and cueing mitosis. The mechanism involved in orienting the mitotic spindle does not appear to sense relative cell stress directly. Instead, division is best predicted by an axis of shape defined by TCJs and is dependent on functional cadherin and the recruitment of LGN. In contrast to this shape-based mechanism, we find that cells may directly sense mechanical stress as a cue for mitotic entry, in a myosin II-dependent manner.

## Author Contributions

**Conceptualization**, S.W. and O.E.J.; **Methodology**, A.N-B., S.W., and O.E.J.; **Software**, A.N.B. **Investigation**, G.A.S-V., G.K.G, T.S., A.N-B. and S.W.; **Formal Analysis**, G.A.S-V., G.K.G. and A.N-B. **Writing – Original Draft**, S.W. and A.N-B., G.A.S-V. and G.K.G; **Writing – Review & Editing**, G.A.S-V., G.K.G., S.W., A.N-B and O.E.J.; **Funding Acquisition**, S.W. and O.E.J.; **Supervision**, S.W. and O.E.J.

## Acknowledgements

ANB was supported by a BBSRC studentship. SW, GSV and GG were supported by a Wellcome Trust/Royal Society Sir Henry Dale Fellowship to SW [098390/Z/12/Z] with additional funding from the Wellcome Trust ISSF [105610/Z/14/Z]. The Bioimaging Facility microscopes used in this study were purchased with grants from BBSRC, Wellcome and the University of Manchester Strategic Fund. We are grateful to Egor Zindy for his assistance in developing in-house nuclei tracking scripts. Thanks to Peter March and Roger Meadows for their help with the microscopy and to Lance Davidson for sharing Cadherin constructs. Also, special thanks to Viki Allan, Tom Millard and Nancy Papalopulu for their critical reading of the manuscript.

## Materials and Methods

### *Xenopus laevis* embryos and microinjection

*Xenopus laevis* embryos were obtained and injected as described previously (Woolner and Papalopulu, 2012). RNA was synthesised as described previously (Sokac et al., 2003) and microinjected at the following needle concentrations: 0.5 mg/ml GFP-α-tubulin; 0.1 mg/ml cherry-histone2B(Kanda et al., 1998); 0.125 mg/ml cadherin 3a full length:6x myc-tag; 0.125 mg/ml cadherin 3a deleted cytosolic domain:6x myc-tag (Kurth et al., 1999). For mosaic expression of β-catenin-GFP (Addgene plasmid #16839, Randall Moon) and GFP-LGN (sub-cloned into pCS2+ from Addgene plasmid #37360, Iain Cheeseman), RNA was injected into a single cell at the 4-cell stage at 0.25 mg/ml (needle concentration). Morpholinos prepared as 1mM stocks (diluted in water) were heated at 65°C for 5 minutes and microinjected at a needle concentration of 1mM and needle volume of 2.5nl into all cells of four-cell stage embryos. The morpholinos used were MHC-B (Myosin Heavy Chain-B, myosin II) MO (5’-CTTCCTGCCCTGGTCTCTGTGACAT-3’; (Skoglund et al., 2008) and standard control MO (5’-CCTCTTACCTCAGTTACAATTTATA-3’; Gene Tools LLC). All embryos were incubated at 16°C for approximately 20 hours prior to animal cap dissection.

### Animal cap dissection and culture

Animal cap tissue was dissected from the embryo at stage 10 of development (early gastrula stage) following a previously described protocol (Joshi and Davidson, 2010), and cultured in Danilchik’s for Amy explant culture media (DFA; 53mM NaCl_2_, 5mM Na_2_CO_3_, 4.5mM Potassium gluconate, 32mM Sodium gluconate, 1mM CaCl_2_, 1mM MgSO_4_) on a 20mm × 20mm elastomeric PDMS (Sylgard 184, SLS) membrane made in a custom mold and coated with fibronectin (fibronectin from bovine plasma, Sigma). Explants were held in place by a coverslip fragment. Each membrane was then incubated at 18°C for at least 2 hours prior to imaging.

### Animal cap stretch manipulation and confocal imaging

Each PDMS membrane was attached to a stretch apparatus (custom made by Deben UK Limited) fixed securely to the stage of a Leica TCS SP5 AOBS upright confocal and a 0.5mm (to remove sag on the membrane) or 8.6mm uniaxial stretch was applied for unstretched and stretched samples respectively. Images were collected on a Leica TCS SP5 AOBS upright confocal using a 20x/0.50 HCX Apo U-V-I (W (Dipping Lens)) objective and 2x confocal zoom. The distance between optical sections was maintained at 5μm and the time interval between each frame was 20 seconds, with each sample being imaged for up to 2.5 hours. For quantification of β-catenin-GFP and GFP-LGN localisation, animal caps were prepared as described but timelapse movies were collected with 2µm optical sections and a time interval of 1 min between frames. Maximum intensity projections of these 3D stacks are shown in the results; except for the GFP-LGN timelapse (Figure 4G), which is an average intensity projection.

### Scanning EM

Uninjected embryos were allowed to develop to stage 10 at 16°C and then animal cap tissue was dissected and allowed to adhere to a fibronectin PDMS membrane as described previously. After 2 hours the animal caps were fixed following a protocol previously detailed by (L. A. Jones et al., 2014). Briefly, the animal caps were fixed in 3.7% PFA and 2.5% Glutaraldehyde in BRB80 buffer (80mM PIPES, 1mM MgCl_2_, 1mM EGTA, pH 6.8) overnight at 4°C. Samples were processed using a high density staining method detailed in full by (Williams et al., 2011) (supplementary protocol), but briefly comprising a 1 hour fix in 2% (wt/vol) osmium tetroxide and 1.5% (wt/vol) potassium ferrocyanide in 0.1M cacodylate buffer. This was followed by a 20 minute incubation in 1% (wt/vol) thiocarbohydrazide and a 30 minute incubation in 2% (wt/vol) osmium teroxide, followed by a final incubation in 1% (wt/vol) uranyl acetate overnight at 4°C. Samples were then stained with freshly prepared Walton’s lead aspartate (0.02M lead nitrate and 0.03M in aspartic acid, adjusted to pH 5.5) for 30 minutes at 60°C prior to dehydration, embedding in Epon 812 (hard formulation), and trimming on a standard microtome. Samples were visualised using a microtome (3View; Gatan) within a Quanta 250 FEG; FEI scanning electron microscope using the following imaging conditions: indicated quadrant magnification of 1600x, accelerating voltage of 3.8kV, pressure at 0.33 Torr. Images were collected at 4000 × 5000 pixels with a dwell time of 10 μs. Raw data was converted to an MRC file stack using IMOD (Kremer et al., 1996; Starborg et al., 2013) and further processed using Imaris software (Bitplane).

### Image analysis

Image analysis was performed using ImageJ (Schneider et al., 2012). Cell division orientation was quantified using the straight-line tool to draw a line between the dividing nuclei of a cell in late anaphase (a stage in mitosis where division orientation is set and the spindle undergoes no further rotation (Woolner et al., 2008; Woolner and Papalopulu, 2012)). Using the ROI manager the angle of division relative to stretch (horizontal axis) was recorded along with the frame and location of the division. Single cell edges and junctions were manually traced 40s before NEB using the freehand paintbrush tool. The whole population of cells in the apical layer of the animal cap was manually traced, along with peripheral junctions and cell centres, using the freehand paintbrush tool. Segmentation of the cell boundaries was performed using in-house Python scripts implementing a watershed algorithm. Geometric features of the cells, such as area and perimeter, were extracted and analysed in Python; for further details on how cell shape was characterised using the segmented images, please see the Supplementary Document. To quantify β-catenin-GFP and GFP-LGN localisation at TCJs in mitotic cells, movies of unstretched and stretched animal caps were analysed as follows: mitotic cells which had non-expressing neighbours were selected at early metaphase. A single optical slice which was level with the centre of metaphase nuclei (visualised by mCherry-H2B) was selected and ROI’s were drawn around TCJs and the corresponding cell edges in Image J. Mean grey values were measured for each ROI and TCJ and cell edge grey values were averaged (mean) for each cell. A ratio between average TCJ intensity and average cell edge intensity (Mean TCJ intensity/Mean cell edge intensity) for each mitotic cell was then calculated.

### Data analysis

The data analysis and plotting was carried out using in-house Python scripts. Statistical tests were performed using the SciPy library (E. Jones and Oliphant E, 2001) and Prism 7 (GraphPad Software, Inc). Mann-Whitney U tests were used to assess if rose histograms were distributed closer to zero. Kolmogorov-Smirnov tests were used to assess if two distributions were significantly different. Otherwise, bootstrapping with 95% confidence intervals, which allow the precision of the estimate to be seen (Nakagawa and Cuthill, 2007), were used to assess significance.

### Immunofluorescence

Embryos were fixed at stage 12 following the protocol previously detailed by Jones et al., (2014)(L. A. Jones et al., 2014). Embryos were incubated in primary and secondary antibodies in TBSN/BSA (Tris-buffered saline: 155mM NaCl, 10mM Tris-Cl [pH 7.4]; 0.1% Nonidet P-40; 10 mg/ml BSA) overnight at 4°C, with five 1 hour washes with TBSN/BSA following each incubation. Primary antibodies were: anti-β-catenin at 1:200 dilution, raised in rabbit (Abcam) and anti c-myc 9E10 at 1:1000 dilution, raised in mouse (Santa-cruz). Alexa Fluor secondary antibodies, anti-rabbit 488 and anti-mouse 568 (Invitrogen) were used at a dilution of 1:400. After staining, embryos were methanol dehydrated, then cleared and mounted in Murray’s Clear (2:1, benzyl benzoate:benzyl alcohol; (Klymkowsky and Hanken, 1991)). Images were collected on a Leica TCS SP5 AOBS inverted confocal using a 63x HCX PL APO (Oil λBL) objective and 1024 × 1024 format. Single confocal slices are shown in the results.

### Implementation of microtubule length division model by Minc et al. (2011)

Details of the model and implementation are given in the Supplementary Document. The predicted torques and corresponding division axes were calculated using in-house Python scripts that are available upon request.

### Implementation of the vertex-based model

The numerical simulations of the vertex-based model were carried out using the same scripts outlined in section 3.8 of (Nestor-Bergmann et al., 2017). Model parameters used for all simulations were (Λ, Γ) = (−0.259, 0.172), determined using a fitting procedure described in (Nestor-Bergmann et al., 2017).

